# Same trait, different genes: pelvic spine loss in three brook stickleback populations in Alberta, Canada

**DOI:** 10.1101/2024.06.27.600703

**Authors:** Jonathan A. Mee, Carolyn Ly, Grace C. Pigott

## Abstract

The genetic basis of phenotypic or adaptive parallelism can reveal much about constraints on evolution. This study investigated the genetic basis of a canonically parallel trait: pelvic spine reduction in sticklebacks. Pelvic reduction has a highly parallel genetic basis in threespine stickleback in populations around the world, always involving a deletion of the pel1 enhancer of *Pitx1*. We conducted a genome-wide association study to investigate the genetic basis of pelvic spine reduction in three populations of brook stickleback in Alberta, Canada. Pelvic reduction did not involve *Pitx1* in any of the three populations. Instead, pelvic reduction in one population involved a mutation in an exon of *Tbx4*, and it involved a mutation in an intron of *Lmbr1* in the other two populations. Hence, the parallel phenotypic evolution of pelvic spine reduction across stickleback genera, and among brook stickleback populations, has a non-parallel genetic basis. This suggests that there may be redundancy in the genetic basis of this adaptive polymorphism, but it is not clear whether a lack of parallelism indicates a lack of constraint on the evolution of this adaptive trait. Whether the different pleiotropic effects of different mutations have different fitness consequences, or whether certain pelvic reduction mutations confer specific benefits in certain environments, remains to be determined.

**Lay Summary:** In this study, we looked for the genetic basis of a well-studied trait in stickleback fish: the pelvic spines. This structure (i.e. the pelvic girdle and attached spines) has a shared developmental basis (and is homologous to) the pelvic bones and hind limbs of all tetrapods (including humans). We know from studying mice, fish, humans, and even manatees that there are several genes that could affect the development of pelvic spines and hind limbs. In one species of stickleback, the threespine stickleback, however, a single gene called *Pitx1* is always involved in the loss of pelvic spines in populations that have adapted to freshwater lakes. This replicated evolution of the same trait in the same environmental conditions is called parallel evolution. It’s remarkable that *Pitx1* is always the gene underlying this adaptive loss of spines in freshwater threespine stickleback populations. We were interested in whether this “genetic parallelism” extended to other species of stickleback that have also evolved the loss of pelvic spines. We looked at three populations of brook stickleback (which are never found in the ocean), each of which contains individuals with and without pelvic spines. We found that the *Pitx1* genetic parallelism does not extend to brook stickleback, and, in fact, the genetic basis of pelvic spine loss differs between populations. In Muir Lake and Astotoin Lake, pelvic spine loss results from a mutation in the *Lmbr1* gene, and in Shunda Lake, pelvic spine loss results from a mutation in the *Tbx4* gene.

## Introduction

Parallel phenotypic evolution is an important observation for inferences about adaptation and the repeatability of evolution. Parallel phenotypic change does not, however, necessarily involve parallel genetic change, and the same phenotype may be caused by different mutations. Given phenotypic parallelism (or adaptive parallelism), genetic parallelism could, conceptually, entail parallel involvement of the same developmental network, the same biological pathway, the same gene (or a paralog), or the same basepair change (James et al. 2023). When (and when not) to expect genetic parallelism is an open question, although there is an intuitive and often-observed tendency for more genetic parallelism when lineages are more closely related (Conte et al. 2012, Blount et al. 2018, Bohutínská & Peichel 2023).

Insights about constraints on evolution can be gained by studying the genetic basis of parallel adaptation. For example, a phenotype that can be produced via a mutation in one of many genes in a biological pathway is less constrained than a phenotype that can only be produced via a mutation in one gene. The genetic basis of a phenotype may also be constrained by evolutionary mechanisms (Yeaman et al. 2018). For example, mutations in different genes may produce the same phenotype but may have different pleiotropic effects, and different mutations might have different expressivity, penetrance, or dominance relationships. As a result, natural selection may impose constraints based on which of the functionally equivalent genetic changes that produce the same phenotype result in the highest fitness. In small and isolated populations, genetic drift may limit genetic diversity or availability of mutations in alternative genes. So, the mutation currently underlying a phenotype in a particular population may be contingent the past availability of one mutation (of several alternative mutations that give in that same phenotype). Overall, one should expect less genetic parallelism between two phenotypically (or adaptively) parallel lineages if any combination of the following is true: the lineages are distantly related, the phenotype has a highly redundant genetic basis, the fitness differences between alternative genetic pathways are small, and/or genetic drift limits the availability of alternative mutations. It is not clear how to generalize expectations regarding genetic parallelism across all combinations of these factors (e.g. the redundancy in biological pathways, the fitness variation among alternative mutations, and the effects of drift), and it is therefore unclear whether the theoretical null expectation should be that genetic parallelism is common or rare. Empirical studies of the genetic basis of parallel phenotypes are therefore central to understanding genetic parallelism and the nature of constraints on evolution.

Sticklebacks (Gasterosteidae) are an established system for studying the genetic basis of phenotypic parallelism. Polymorphism in the presence-absence of pelvic spines and pelvic girdles has been observed among populations in four stickleback genera: *Apeltes*, *Culaea*, *Gasterosteus*, and *Pungitius*. In *Gasterosteus aculeatus* and Northern European populations of *Pungitius pungitius*, pelvic spine loss or reduction is caused by a deletion mutation in the pel1 enhancer affecting the expression of *Pitx1* in the pelvic region during development (Shapiro et al. 2004, Shapiro et al. 2006, Coyle et al. 2007, Chan et al. 2010, Klepaker et al. 2013, Shikano et al. 2013, Kragesteen et al. 2018, Thompson et al. 2018, Xie et al. 2019). Hence, there is genetic parallelism among these two stickleback genera that diverged approximately 26 million years ago (Shapiro et al. 2006, Varadharajan et al. 2019). But, pelvic spine reduction among populations of *P. pungitius* in North America does not involve *Pitx1* (Shikano et al. 2013), suggesting that there is redundancy in the genetic basis of this trait. Indeed, among the genes in the regulatory network that influences the development of pelvic structures in fishes (i.e. pelvic spines or fins) or hind limbs in tetrapods, a mutation in any number of different genes (e.g. in *Pitx1*, *Tbx4*, *Wnt8c*, or *Hox9*-*Hox11*) could result in reduction or loss (Swank et al. 2021).

Regardless of the genetic basis of stickleback pelvic spine reduction, there is an abundance of evidence from studies of sticklebacks supporting the assumption that pelvic spine phenotype is influenced by selection, and alternative pelvic morphs (e.g. spined versus unspined) may be under directional or balancing selection within a population depending on the context.

Selection on pelvic spine morphs is influenced by predation (for or against spines depending on the predator) and by the physiological or resource costs of growing bony structures, particularly in low-calcium freshwater environments (Reist 1980a, Reist 1980b, Giles 1983, Bell et al. 1993, Reimchen 1994, Rundle et al. 2003, Lescak et al. 2012, Spence et al. 2013, Miller et al. 2017, Haines et al. 2023).

This study addressed the question, to what extent does pelvic spine reduction or loss involve a parallel genetic basis among stickleback species and populations? This question was addressed by investigating the genetic basis of pelvic spine reduction in brook stickleback (*Culaea inconstans* Kirtland, 1840), and by comparing the findings to the well-studied genetic basis of pelvic spine reduction in threespine and ninespine stickleback involving *Pitx1*, as described above. Three populations of brook stickleback in Alberta, Canada, with different proportions of spined, partially spined (a.k.a. intermediate or vestigial), and unspined individuals were used for this study. The difference in the pattern of polymorphism in these three lakes may be due to a difference in the genetic basis of the pelvic reduction. For example, different mutations (e.g. in different genes) might have different expressivity, penetrance, or dominance relationships that result in different phenotypes in heterozygous individuals, and, as a result, different proportions of partially spined individuals. The three populations of brook stickleback (i.e. from three different lakes) included in this study were selected because the frequencies of pelvic spine morphs differed between populations, and because the polymorphism has persisted in these lakes with relatively high and stable frequencies of pelvic spine morphs for least five decades (Nelson 1977, Klepaker et al. 2013, Lowey et al. 2020). We had a prior expectation that we would find evidence of the involvement of *Pitx1* in pelvic spine polymorphism in brook stickleback. If the molecular features that promote fragility (or mutability) of *Pitx1* enhancer sequences in threespine stickleback (Xie et al. 2019) are conserved across genera, one may even expect a deletion of the pel1 enhancer to be involved in brook stickleback pelvic spine reduction. We had no prior expectation regarding the likelihood of parallelism or non-parallelism in the genetic basis of pelvic spine loss among different brook stickleback populations.

## Materials and Methods

### Sample collection

Brook Stickleback were collected from Muir Lake and Shunda Lake in 2017 and 2019, and from Astotin Lake in 2022 (Fig 1). Astotin Lake and Muir Lake are comprised of a roughly 50% mix of spined and pelvic-reduced brook stickleback (Nelson 1977, Lowey et al. 2020). In Astotin Lake, about 75% of pelvic-reduced individuals completely lack any pelvic structure, and the remainder of the pelvic-reduced individuals have partial pelvic reduction (e.g. missing a spine on the left or right side, or having a pelvic girdle but no spines). In Muir Lake, these intermediate morphs (with partial pelvic reduction) occur at frequency of less than 2% (Lowey et al. 2020). The brook stickleback population in Shunda Lake is comprised mostly of individuals with a complete pelvic girdle and spines, and a minority (about 20%) of individuals in Shunda Lake have some degree of pelvic loss, about 25% of which (i.e. ∼5% of the total population) completely lack any pelvic structure (Lowey et al. 2020). Fish were collected using 0.5cm mesh minnow traps set along lake margins within 5m of shore at a depth of up to 2m. All traps were retrieved within one to twelve hours after being set. Brook stickleback were anesthetized and euthanized in an overdose mixture of lake water and eugenol, caudal fins were preserved in 95% ethanol, and all other tissues were preserved in 70% EtOH. Sampling permits were issued by the Government of Alberta and Parks Canada, and fish handling protocols were approved by the Animal Care Committee at Mount Royal University (Animal Care Protocol ID 101029 and 101795).

**Figure 1.**
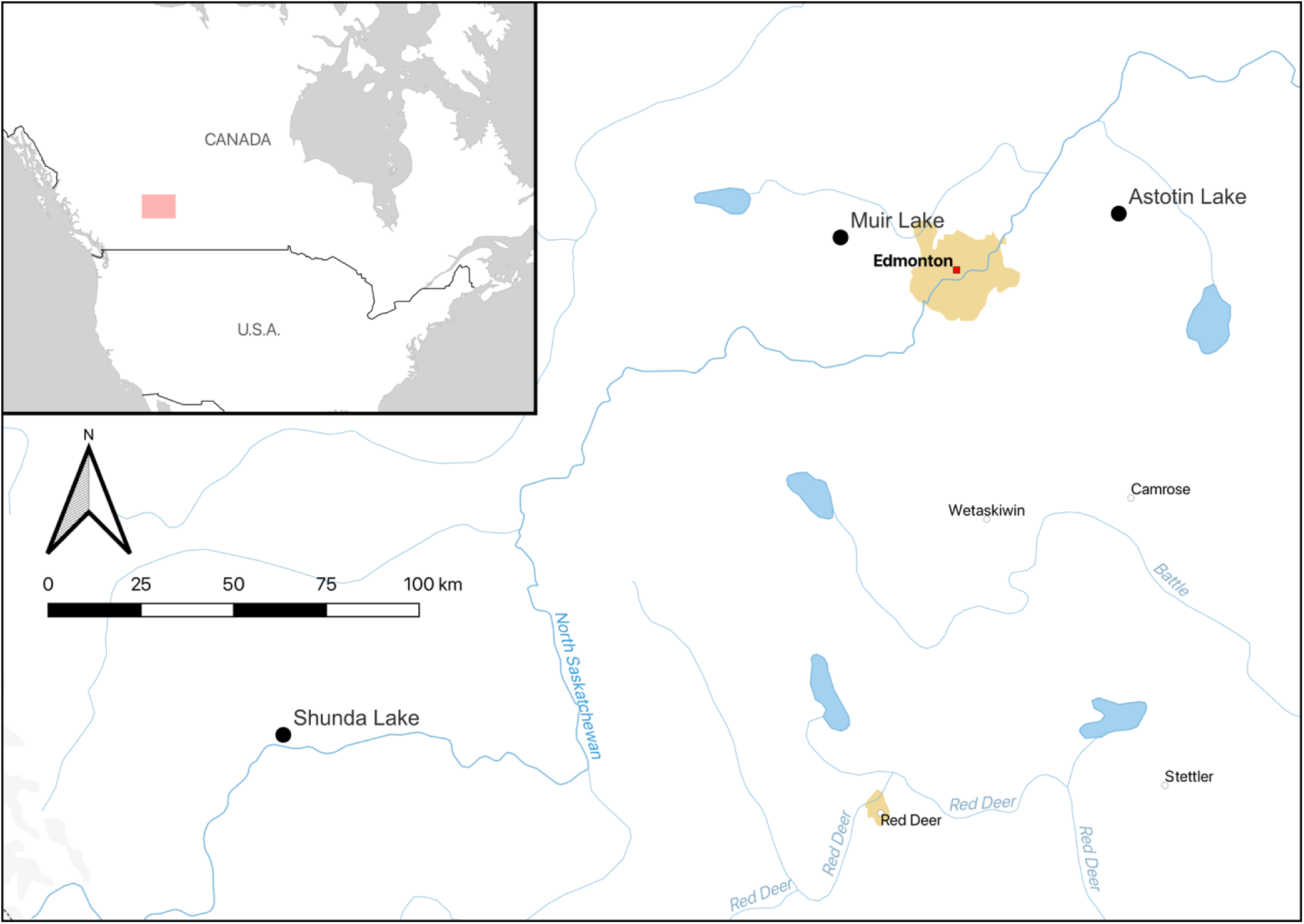
Locations of the three populations of brook stickleback sampled for this study (Astotin Lake, Muir Lake, and Shunda Lake). The pink box in the inset map shows the area displayed in the main map. The map was made with QGIS using Natural Earth Data.

### Pelvic phenotypes

Spined and unspined individuals were initially identified at the site of capture based on close visual inspection and articulation of pelvic spines with fine-tipped tweezers. Sex was also evaluated at the site of capture by examining gonads and by noting the presence of nuptial colouration. Subsequently, brook stickleback specimens were bleached and dyed using alizarin red to clearly visualize calcium-containing oocytes (such as those in pelvic bones) following a protocol adapted from Xie et al. (2019). After bleaching and staining, photographs of each specimen were taken using a Nikon SMZ-745T Zoom Stereo Photo Microscope. Photos of each specimen, focusing on the entire pelvic area, the left pelvic girdle, the right pelvic girdle, the left pelvic spine, and the right pelvic spine, including a scale bar within the focal area for each photo, were used to verify the pelvic phenotype for each individual, and each individual was scored as pelvic-present, pelvic-absent, or intermediate (as per Klepaker et al. 2013; Figure S1). In addition, measurements of the length of both pelvic spines (when present) and the area of the left and right pelvic girdle (when present) were recorded using Nikon NIS-Elements D software.

Because left-bias in individuals with pelvic vestiges has been identified as a hallmark of *Pitx1* mutation (Bell et al. 2007, Alvarado et al. 2011), we used a standard measure of asymmetry from Palmer (1994) to further characterize pelvic phenotype for each individual: (L − R)/[(L + R)/2)].

### Whole genome sequencing

Genomic DNA was extracted from each individual using a Qiagen DNEasy Blood and Tissue kit. Purified DNA samples with a DNA concentration > 1 ng / uL (measured with a Qubit Fluorometer and a Broad Range DNA assay kit) and with a 260/280 nm absorbance ratio > 2 (measured with a Nanodrop Spectrophotometer) were sent to Genome Québec for shotgun DNA library preparation and paired-end (150bp) sequencing on an Illumina HiSeqX or NovaSeq6000 with an S4 flow cell. In 2017, 40 individuals from Muir Lake and 40 individuals from Shunda Lake were sequenced on four HiSeqX lanes. An additional 58 individuals from Muir Lake and 57 individuals from Shunda Lake (from samples collected in 2017 and 2019) were sequenced on one lane of NovaSeq6000 S4. In 2022, 120 individuals from Astotin Lake were sequenced on one lane of NovaSeq6000 S4. All raw data was deposited on the Short Read Archive (SRA) database: accession numbers PRJNA895038 (Astotin Lake), PRJNA838068 (Muir Lake), and PRJNA838194 (Shunda Lake).

### SNP calling

To balance SNP quality (i.e. a low false positive rate) and processing time (Jasper et al. 2022), we chose to use the following pipeline employing BCFtools to call SNPs (Danecek et al. 2021). Quality filtering (phred quality >= Q15, unqualified bases <=40%), length filtering (> 15bp), adapter trimming, and, for NovaSeq reads, polyG tail trimming (minimum length = 10) were conducted using fastp v.0.20.1 (Chen et al. 2018). No reference genome for *C. inconstans* was available, and the *Pungitius pungitius* fPunPun2.1 (GCF_949316345.1) reference genome assembly (Varadharajan et al. 2019) was used for alignment using bwa v.0.7.17 (Li 2013).

Samtools v.1.17 (Danecek et al. 2021) was used to generate, sort, and index BAM files, skipping alignments with map quality < 10 (-q 10). Duplicates were then marked and removed using MarkDuplicates in picard v.2.26.3 (http://broadinstitute.github.io/picard/). The BAM files were then reindexed using samtools, and the cleaned and indexed BAM files were realigned by running RealignerTargetCreator and IndelRealigner from the Genome Analysis Toolkit v.3.8 (DePristo 2011). To identify single nucleotide polymorphisms (SNPs), genotype likelihoods were generated by running mpileup, specifying a minimum mapping quality of 5 for retention of an aligned variant, then generated SNP VCF files by running call in BCFtools v.1.11 (Danecek et al. 2021). Using VCFtools (Danecek et al. 2021), the individual VCF files from each population were combined into a single population VCF file using the vcf-concat command then sites with quality value below 30 (--minQ 30), with genotype quality below 20 (--minGQ 20), with read depth below 5 (--minDP 5), and with more than 2 alleles (--max-alleles 2) were removed. The full SNP-calling pipeline is available at https://github.com/jon-mee/culaea_wgs_SNPs.

### Genome-wide genotype-phenotype association

To analyze genome-wide genotype-phenotype association, and for other downstream analyses, PLINK v.1.90b4.6 (Purcell et al. 2007) was used to create a filtered subset with a minimum allele frequency of 1% (--maf 0.01) and a maximum of 20% missing data per SNP (-- geno 0.2). PLINK was then used to create a linkage disequilibrium pruned (LD-pruned) set of SNPs by removing SNPs within 50 kb windows (at 5kb steps) that were above a variance threshold factor of 2. GCTA (Yang et al. 2011) was used to estimate a genetic relatedness matrix (GRM) from the LD-pruned SNPs. Finally, to analyze genome-wide association between SNPs and pelvic phenotypes, GCTA was used to perform a mixed linear model association analysis (MLMA) using the full unpruned filtered SNP set using the GRM to control for population stratification. To visualize the results of the MLMA, the QQMAN package (Turner 2014) was used in R (R Core Team 2018). Genes associated with outliers (using a significance threshold of p < 10^-7^) in the MLMA analysis were retrieved from the *P. pungitius* genome annotation: https://www.ncbi.nlm.nih.gov/gdv/browser/genome/?id=GCF_949316345.1. The vcfR package (Knaus and Grünwald 2017) was used to retrieve SNP genotypes within a region +/- 100Kb from the most significant outlier (or focal) SNPs, and the Pearson correlation (r^2^) between each SNP within the outlier region and the focal SNP to estimate LD using the cor function were calculated in R (R Core Team 2022).

### Genetic diversity and selection metrics

To characterize patterns of population genetic diversity, the snpR package in R (Hemstrom and Jones, 2022) was used to calculate π, H_E_, F_IS_, and pairwise F_ST_. Tajima’s D was calculated in 1000bp windows for all three populations using the SNP data filtered for quality and depth of coverage, but without filtering for minimum allele frequency (so as to not bias Tajima’s D estimates). Finally, to evaluate the genetic structure within and between each population, and to determine whether any within-population genetic structure is associated with pelvic spine polymorphism, ancestry models were constructed using ADMIXTURE v.1.3.0 (Alexander et al. 2009) with the number of ancestral populations (K) set to 1 through 6, and a principal component analysis (PCA) was conducted using PLINK v.1.90b4.6 (Purcell et al. 2007).

## Results

The numbers of spined, intermediate, and unspined individuals included in the analyses for each lake are listed in Table 1. An attempt was made to include a balanced sample of males and females for each phenotype from each lake (Table 1), but sex was not included as a variable in the analyses. Before filtering for minimum allele frequency and missing data per SNP, there were 20,706,142 SNPs discovered in Astotin Lake, 20,531,270 SNPs in Muir Lake, and 20,407,490 SNPs in Shunda Lake. After filtering, 1,250,805 SNPs were retained in Astotin Lake, 1,327,906 SNPs in Muir Lake, and 1,166,516 SNPs in Shunda Lake. The LD-pruned datasets (for GRM estimation) retained 478,612 SNPs in Astotin Lake, 424,489 SNPs in Muir Lake, and 381,815 SNPs in Shunda Lake.

**Table 1.** Number of brook stickleback sampled for whole-genome sequencing and used in analyses of genome-wide genotype-phenotype association for this study.

|  | Pelvic Phenotype |  |  |
| --- | --- | --- | --- |
|  | Spined | Intermediate | Unspined |
| Astotin Lake |  |  |  |
| female | 32 | 13 | 30 |
| male | 17 | 1 | 9 |
| unknown sex | 11 | 1 | 5 |
| Muir Lake |  |  |  |
| female | 18 | 1 | 19 |
| male | 29 | 1 | 30 |
| Shunda Lake |  |  |  |
| female | 21 | 19 | 9 |
| male | 28 | 14 | 6 |

A GWAS was performed for each lake separately based on the filtered (not LD-pruned) datasets and using the GRM estimation from the LD-pruned datasets. In Astotin Lake and Muir Lake, there was a single clear peak of association on chromosome 19 (Figure 2a and 2b). The SNP with the strongest association on chromosome 19 was the same in both lakes (position 4,425,247), and was located within the fifth intron of the limb development membrane protein 1 (*Lmbr1*) gene (https://www.ncbi.nlm.nih.gov/gene/119198937), which contains a conserved regulatory element that drives the spatio-temporal expression of the *sonic hedgehog* (*Shh*) gene (Park et al. 2008). In Shunda Lake, the SNP with the strongest association was on chromosome 3 (Figure 2c). This SNP (position 18,698,851 on chromosome 3) was located within an exon of the T-box transcription factor 4 (*Tbx4*) gene (https://www.ncbi.nlm.nih.gov/gene/119214652).

**Figure 2.**
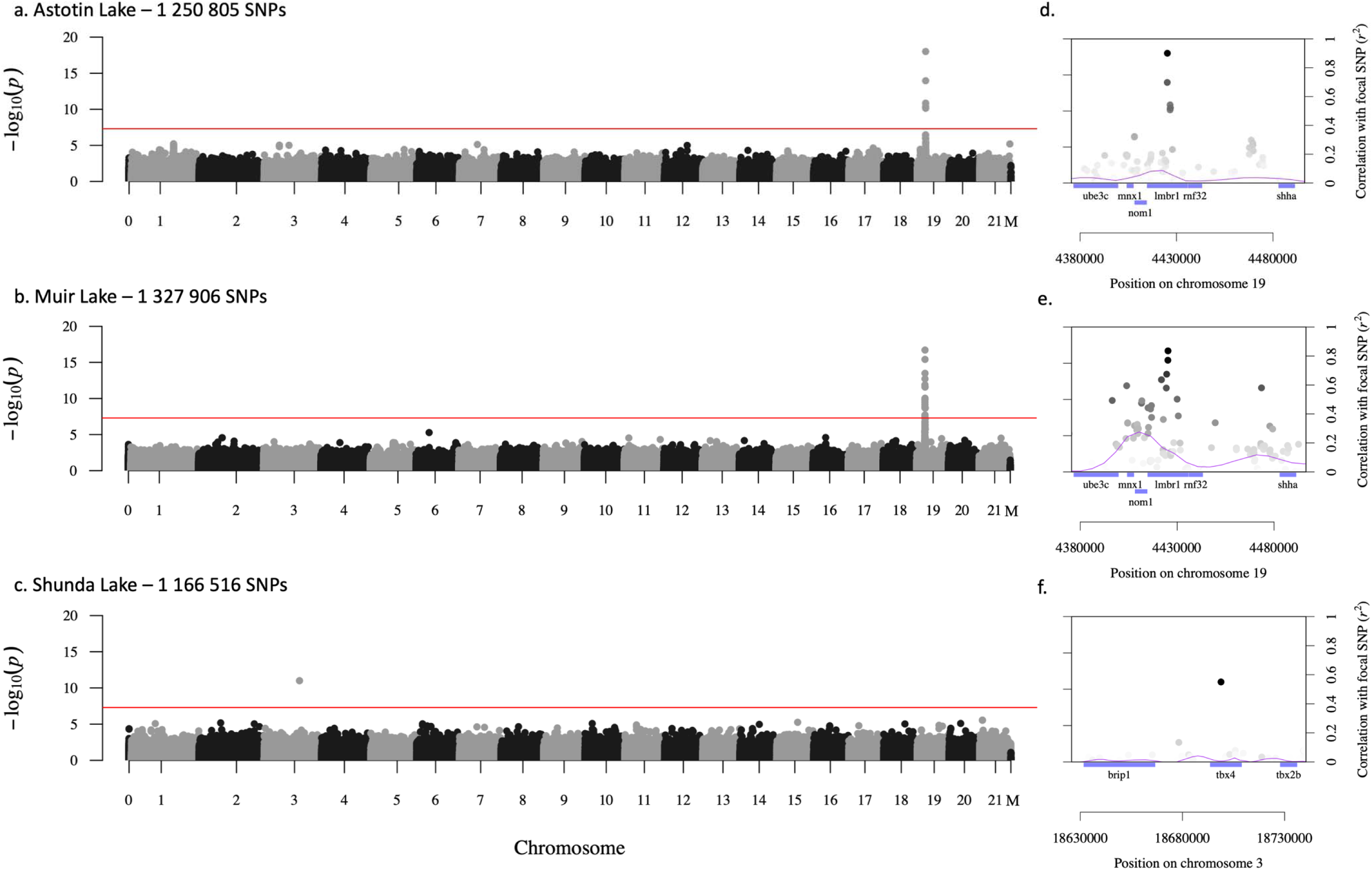
GWAS results for a) Astotin Lake, b) Muir Lake, and c) Shunda Lake. Chromosome zero (i.e. the zero at the left of each x-axis) captures all SNPs in unassembled scaffolds. The outlier regions for each lake (panels d, e, and f respectively) show the locations of genes identified from the *Pungitius pungitius* genome annotation shown in blue below the axis. Genes are labeled with the gene symbol from the annotation. The purple line shows a Lowess-smoothed curve of linkage disequilibrium across the outlier region (measured as correlation with the focal SNP; r^2^ on the right-hand axis). Shading of points in the right-hand panels corresponds to the magnitude of correlation with the focal SNP. The left-hand axis in panels d, e, and f is the same as (and aligned with) panels a, b, and c, respectively.

As expected, there was a high proportion of intermediate pelvic phenotypes in Shunda Lake, a moderate proportion of intermediate phenotypes in Astotin Lake, and a low proportion of intermediate pelvic phenotypes in Muir Lake (Table 1). The majority of intermediate phenotypes were heterozygous in Astotin Lake, whereas the majority of intermediate phenotypes were homozygous for the pelvic-loss allele in Shunda Lake. To visualize patterns of dominance, penetrance, and expressivity for the genes underlying pelvic polymorphism, a generalized linear model with a binomial distribution and a logit link function was conducted in R (R Core Team 2022) to fit a logistic regression between the pelvic phenotype and the focal SNP genotype for each individual (Figure 3). The *Lmbr1* pelvic reduction allele was more dominant in its phenotypic effect than the *Tbx4* pelvic reduction allele. By treating pelvic phenotype as a quantitative trait, and assigning values of 1 and 0 for spined and unspined, respectively, and a value of 0.5 for intermediate phenotypes, the dominance effect of the pelvic reduction allele in each population was calculated as *d* = (*x*_TT_ – *x*_TC_)/(*x*_TT_ – *x*_CC_), where *x_ii_* is the mean phenotype for each genotype. For *Lmbr1*, *d* = 0.9112 in Astotin Lake and *d* = 0.9474 in Muir Lake. For *Tbx4*, *d* = 0.1089 in Shunda Lake. Penetrance (i.e. the proportion of homozygous mutants that have pelvic reduction) was 100% for *Lmbr1* and 97.78% for *Tbx4*. Expressivity was relatively high for *Lmbr1*: 85.2% of pelvic-reduced individuals (unspined or intermediate) are unspined, and the remainder are intermediate. Expressivity was, in contrast, low for *Tbx4*: only 25.5% of pelvic-reduced individuals (unspined or intermediate) are unspined.

**Figure 3.**
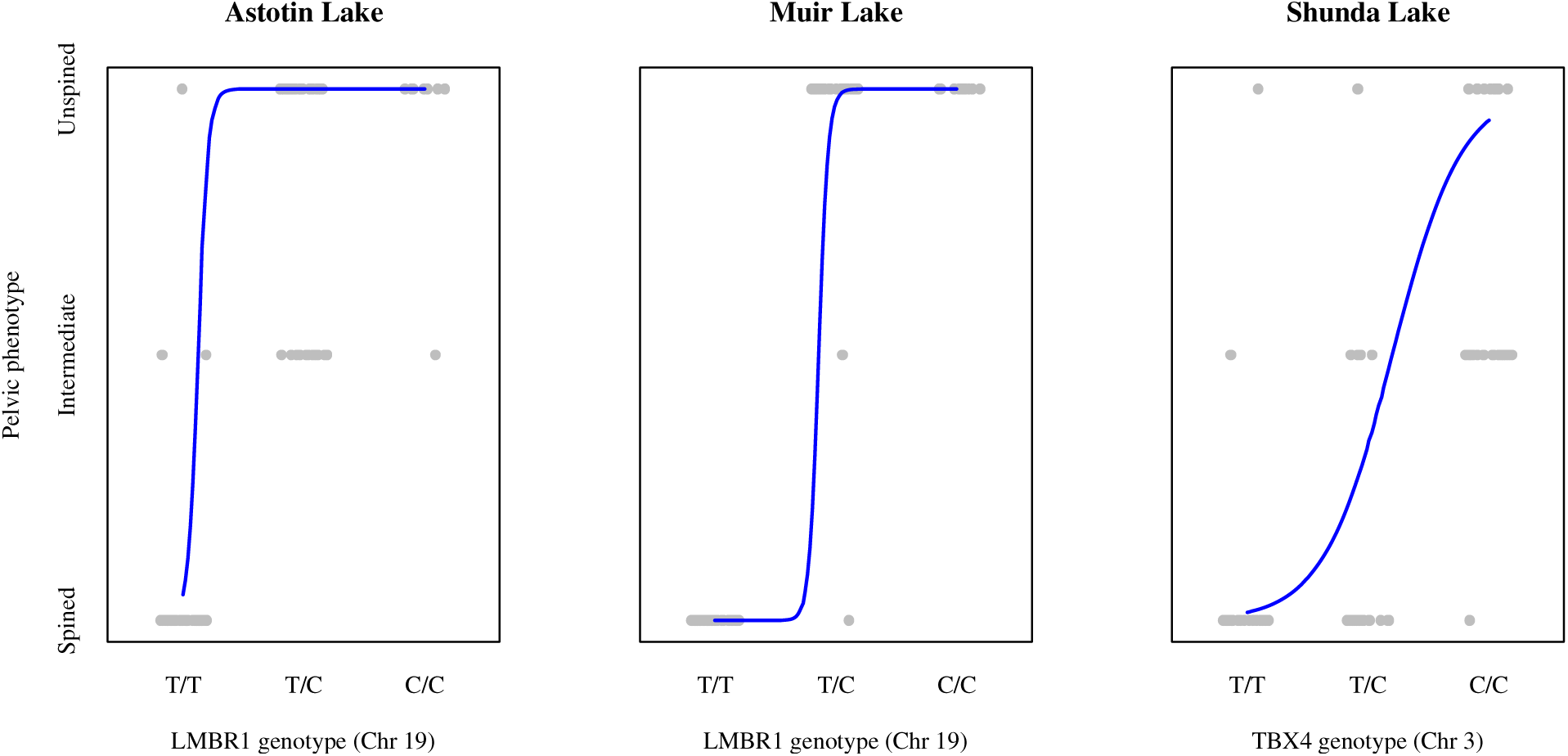
Different patterns of dominance, penetrance, and expressivity of pelvic-reduction mutations in different populations. Each grey point shows the genotype and phenotype for one individual (with random jitter to space out the points on the horizontal axis). The lines show the fit of a logistic regression between pelvic phenotype and genotype for each lake.

Patterns of pelvic asymmetry differed between lakes with different mutations.

Heterozygotes for the *Lmbr1* pelvic-reduction allele (in Astotin Lake and Muir Lake) tended to have right-biased pelvic spines and pelvic girdles, whereas heterozygotes for the *Tbx4* pelvic- reduction allele (in Shunda Lake) tended to have slightly left-biased spines but no bias in pelvic girdle reduction (Figure 4). We used the CAR package in R (Fox & Weisberg 2019) to conduct two-way ANOVAs with type II sums of squares to test the significance of these patterns of asymmetry in pelvic spine length and pelvic girdle area. We tested for differences between genes (*Lmbr1* versus *Tbx4*), for differences between genotypes (non-mutants versus heterozygotes), and for interaction effects. We found a significant interaction effect between gene and genotype for pelvic spine length (F_1,115_ = 7.9132, p = 0.006), but not for pelvic girdle area (F_1,118_ = 3.7414, p = 0.055). None of the main effects (i.e. of gene or genotype) were significant.

**Figure 4.**
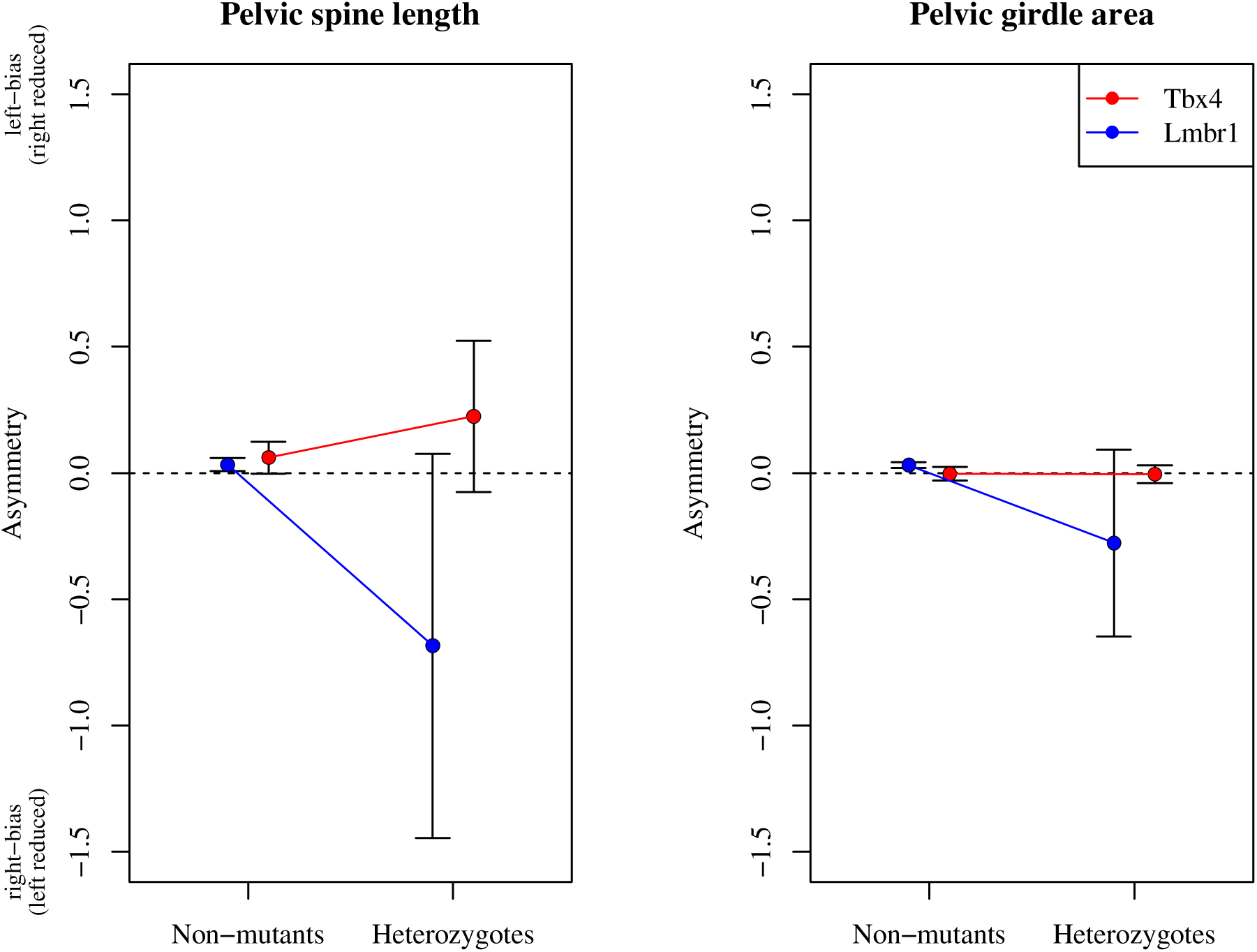
Asymmetry in pelvic spines in non-mutant individuals (typically with complete pelvic girdles and two spines) and in heterozygotes (typically with intermediate pelvic morphology). It was not possible to measure asymmetry in homozygous mutants with intermediate pelvic morphology (most had no pelvic structure at all). Error bars show one standard error.

Shunda Lake had the lowest nucleotide diversity (π) and expected heterozygosity (H_E_) of the three lakes in this study (Table 2). All three lakes had strongly negative F_IS_ values, indicating an excess of heterozygotes, and values of Tajima’s D were, on average, positive (Table 2), suggesting an overall trend of balancing selection. The values of Tajima’s D within the 1000bp windows immediately framing the loci associated with pelvic loss were, however, negative, and there was no clear positive or negative trend in the values of Tajima’s D in the genomic regions adjacent to the loci associated with pelvic loss (Figure 5). ADMIXTURE and PCA analyses revealed no evidence for within-lake structure associated with pelvic spine phenotype, but did reveal strong between-lake genetic differentiation (supplementary figures S2-S4).

**Figure 5.**
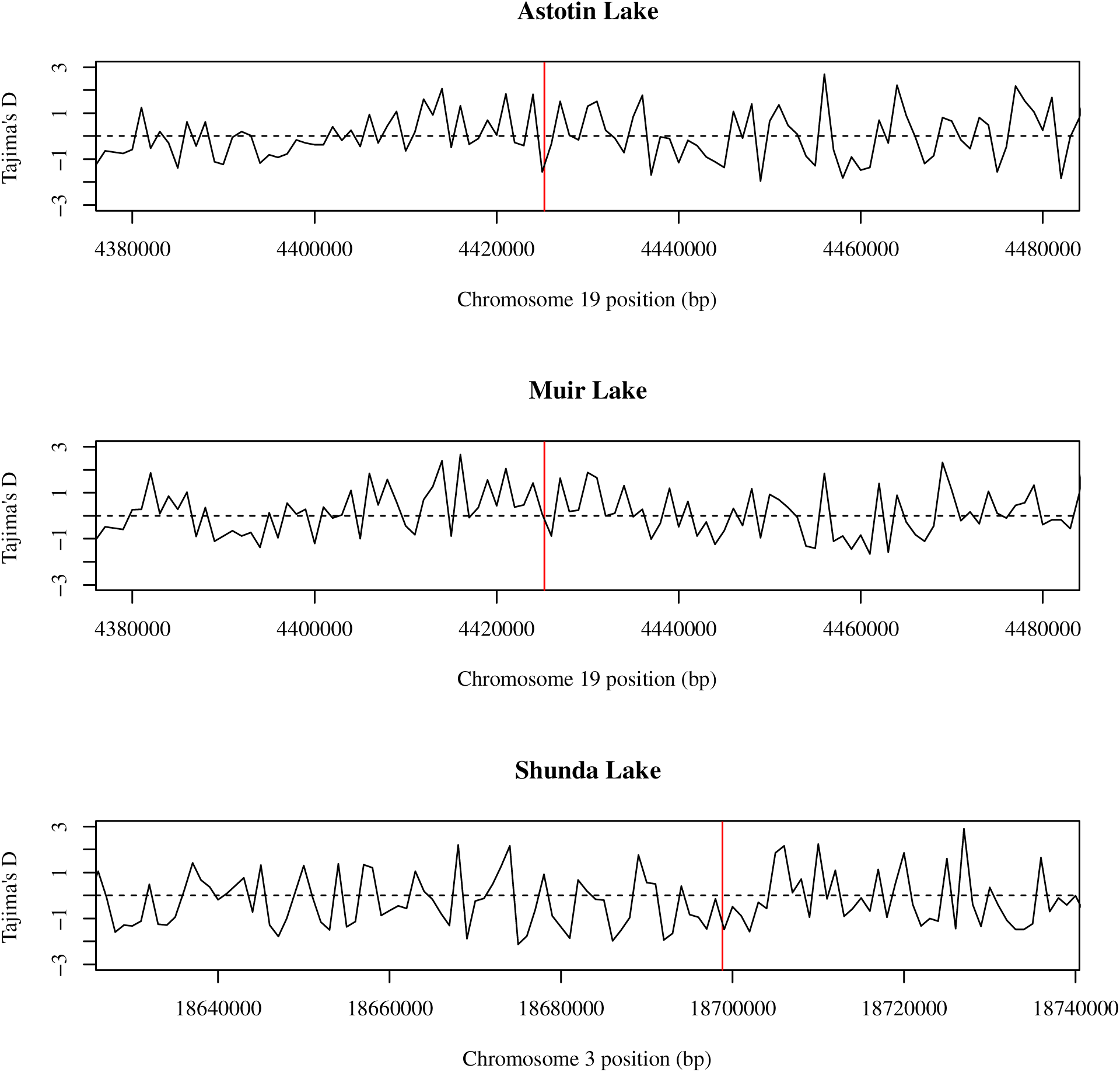
Tajima’s D plotted in 1000bp windows within the outlier regions depicted in Figure 2. The vertical red line shows the location on the horizontal axis of outlier SNP, and the shading denotes the transcribed sequence of the associated gene: *Lmbr1* in Astotin Lake and Muir Lake, *Tbx4* in Shunda Lake.

**Table 2.** Population genetic diversity and selection metrics. H_O_ = observed heterozygosity. H_E_ = expected heterozygosity. π = nucleotide diversity. F_IS_ = inbreeding coefficient. D = Tajima’s D.

| | $H_O$ | $H_E$ | $\pi$ | $F_{IS}$ | Mean $D$ |
| --- | --- | --- | --- | --- | --- |
| Astotin Lake | 0.283 | 0.218 | 0.219 | -0.294 | 0.458 |
| Muir Lake | 0.297 | 0.219 | 0.220 | -0.352 | 0.574 |
| Shunda Lake | 0.295 | 0.208 | 0.209 | -0.416 | 0.506 |

## Discussion

Numerous genes could potentially be involved in pelvic reduction, loss, or malformity in fishes and tetrapods (Swank et al. 2021). Nonetheless, previous studies have frequently, if not consistently, implicated *Pitx1* as the gene underlying reduction or loss in natural populations (Shapiro et al. 2006). For example, *Pitx1* has been implicated in pelvic reduction in manatees and in at least two species of stickleback (Shapiro et al. 2006). In brook stickleback, however, there was no evidence for the involvement of *Pitx1* in pelvic reduction. Instead, two different genes were involved in pelvic spine reduction in different populations. In Astotin Lake and Muir Lake, pelvic reduction involved a mutation in an intron of the *Lmbr1* gene. In Shunda Lake, pelvic reduction involved a mutation in an exon of the *Tbx4* gene. This result suggests a lack of parallelism among species of stickleback and among populations of brook stickleback at the level of the gene, although all these genes likely interact in a single conserved core regulatory network involved in the formation of hindlimbs (Park et al. 2008, Duboc et al. 2021, Swank et al. 2021).

Whether or not this lack of gene-level parallelism should be interpreted as a lack of constraint on the genetic basis of pelvic reduction remains an open question. There could be better and worse ways, in terms of fitness, to achieve pelvic reduction. Assuming that pelvic phenotype is under selection (as ample evidence would suggest – as per the introductory paragraph on this topic, above), it is possible that pelvic polymorphism in these populations is driven by balancing selection on whichever pelvic-reduction mutation happened to arise in each population. Brook stickleback are an entirely North American freshwater species, and most of their populations, especially in Canada, are found in lakes that are isolated from one-another, that were colonized relatively recently following the last glacial maximum roughly 15,000 years ago, and that frequently undergo winter die-backs under ice when lakes become hypoxic. Hence, the likelihood that genetic drift has affected the degree of parallelism among brook stickleback populations is high. It is possible that a universally optimal genetic basis for pelvic spine reduction confers the highest fitness (e.g. has the fewest negative pleiotropic effects), but that some brook stickleback populations are “making do” with whatever mutation is available.

Alternatively, perhaps the best (i.e. highest-fitness) mutation for pelvic spine reduction is dependent on the specific environmental conditions experienced in each population. For example, in the populations in this study, the *Lmbr1* mutation might be the best way to achieve pelvic reduction in Astotin Lake and Muir Lake, whereas the *Tbx4* mutation may be the best way to achieve pelvic reduction in Shunda Lake. These alternatives (i.e. non-parallelism due to drift or due to local adaptation) could be explored by raising fish with different pelvic reduction mutations in a common environment and recording pleiotropic phenotypic effects that might be universally deleterious (e.g. reduced growth rate or fecundity) or that could be adaptive (e.g. effects on trophic morphology).

Brook stickleback are remarkably variable in their morphological and ecological characteristics, which has been noted ever since the species was first given its specific epithet: *inconstans* (Kirtland 1840). Of note, pelvic reduction is correlated with different morphological and ecological consequences in Muir Lake and Shunda Lake (Willerth et al. 2022, Mee et al. 2023). Whether the morphological and ecological differences between lakes, including the different correlates of spine reduction in different lakes, are driven by the different genes underlying spine reduction is unknown. It is possible that different environments in Muir lake and Shunda Lake confer some locally adaptive benefit to *Lmbr1* or *Tbx4* mutations in these lakes. Alternatively, lake-specific differences could be driven by phenotypic plasticity. It does seem likely, however, that the pattern of asymmetry that we observed in intermediate individuals (i.e. in individuals with pelvic vestiges), which differs with *Lmbr1* or *Tbx4* mutations (Figure 4), is an effect of the different mutations in different individuals. Left-bias in individuals with pelvic vestiges has been identified as a hallmark of *Pitx1* mutation (Bell et al. 2007, Alvarado et al. 2011). These results suggest that other mutations underlying pelvic reduction can cause a right-bias (*Lmbr1*) or no bias (*Tbx4*) in pelvic vestiges, which may provide more insight when interpreting patterns of pelvic asymmetry (for example, in fossil data or preserved samples).

The lack of gene-level parallelism in pelvic spine reduction across stickleback species is less surprising than the lack of gene-level parallelism among brook stickleback populations. We expect less genetic parallelism when lineages are less closely related (Conte et al. 2012, Blount et al. 2018, Bohutínská & Peichel 2023). The molecular features that appear to be conserved within threespine stickleback and that cause fragility in the *Pitx1* enhancer, thereby explaining the within-species gene-level parallelism in threespine stickleback (Xie et al. 2019), may not be present in brook stickleback. Identifying the mechanisms underlying patterns of (non)parallelism withing and among species (e.g. molecular features, genetic drift, or local adaptation) would certainly help solidify our understanding of the repeatability of and constraints on evolution.

## Author Contributions

JAM conceived and designed the study, collected and analyzed the data, and wrote the manuscript.

## Acknowledgements

Heartfelt thanks go to all who helped me learn how to call SNPs including Brandon Lind, Sara Smith, Moroni Lopez, Sam Yeaman, Gabriele Nocchi, and Greg Owens. Thanks also to Sam Yeaman for comments on an earlier version of this manuscript. Field work was assisted by undergraduate students Marin Flannagan, Emily Franks, Kaitlyn Willerth, Bryce Carson, and Alyce Straub. Field work support was provided by Melanie Rathburn and Alex Farmer. Lab support was provided by Lindsay Leahul and Ava Zare. Funding for this project was provided by an NSERC Discovery Grant and an MRU Faculty of Science and Technology PCYI award.

## Data Accessibility

The raw sequence data, which includes each individual’s sex and spine phenotype in the metadata, was deposited on the Short Read Archive (SRA) database: accession numbers PRJNA895038 (Astotin Lake), PRJNA838068 (Muir Lake), and PRJNA838194 (Shunda Lake). The SNP analysis pipeline is archived on GitHub at https://github.com/jon-mee/culaea_wgs_SNPs. All other data are archived on Dryad (DOI: 10.5061/dryad.v9s4mw74r). Temporary link for peer review: https://datadryad.org/stash/share/6ZAYHQ_70kPK8kFmYNTNwaejcguGPALWnoXZbSrG358.

---

**Figure S1.**
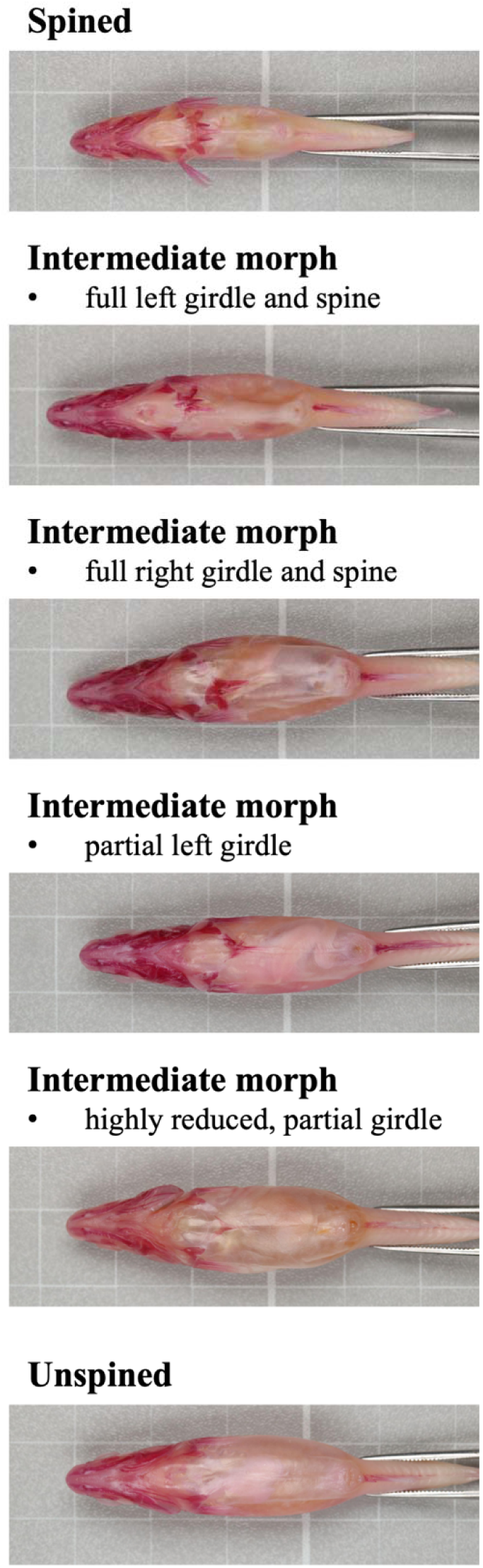
Example brook stickleback pelvic morphologies. All images are of samples from Astotin Lake, Alberta.

**Figure S2.**
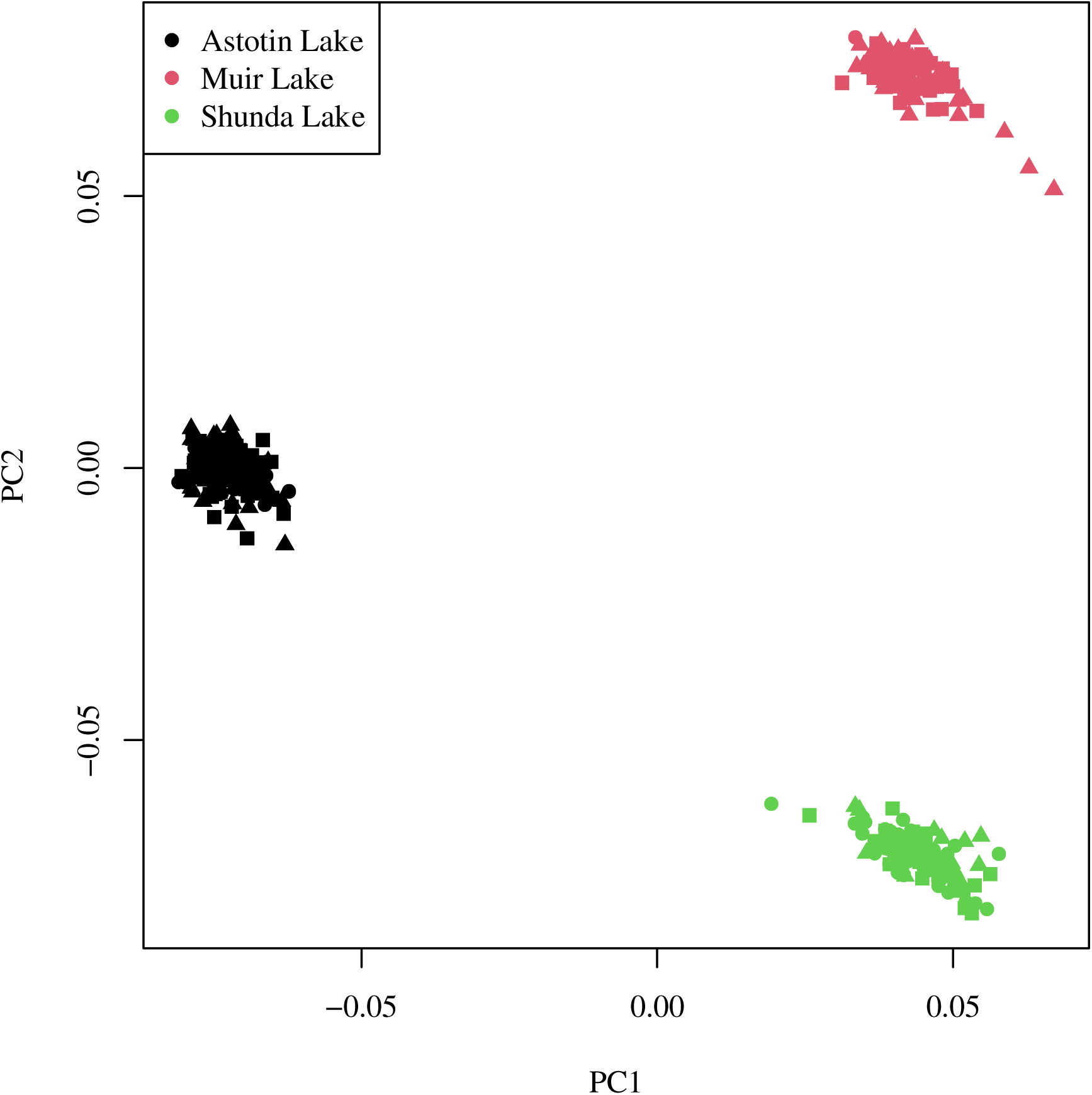
PCA plot based on SNP data from brook stickleback populations in Astotin Lake, Muir Lake, and Shunda Lake. Point shapes denote pelvic spine phenotype: square = spined, circle = intermediate, triangle = unspined.

**Figure S3.**
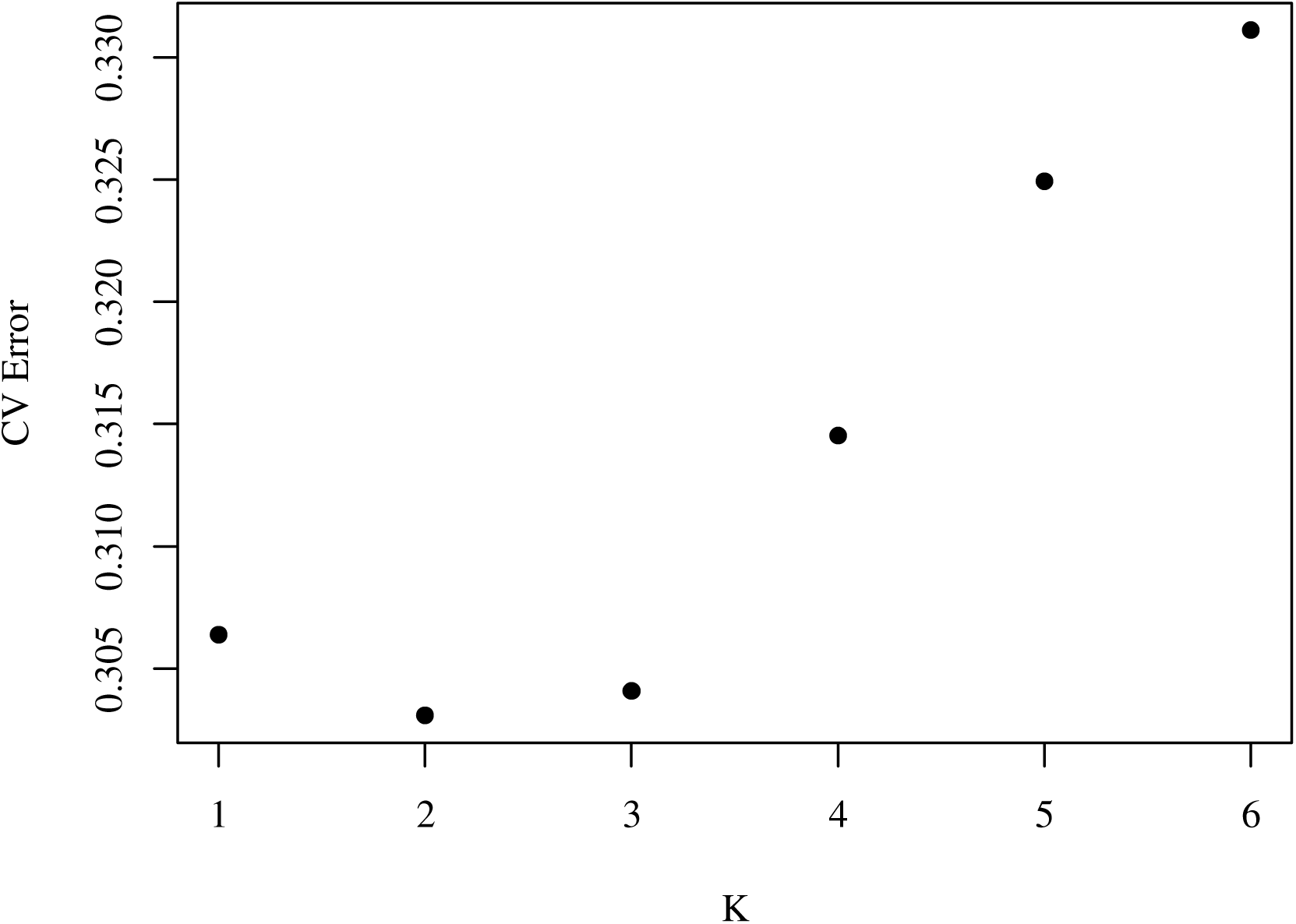
Cross validation error from ADMIXTURE analysis of SNP data from brook stickleback populations in Astotin Lake, Muir Lake, and Shunda Lake, with K set 1 to 6, and 10 replicates of each value of K.

**Figure S4.**
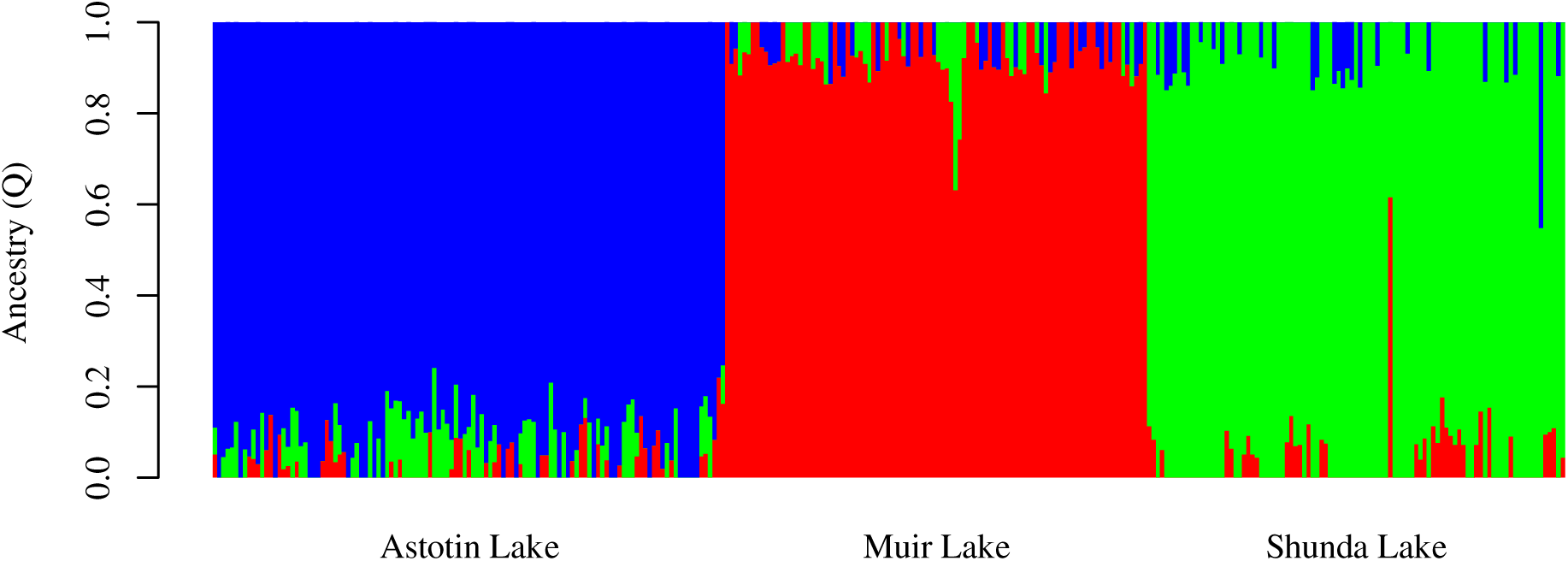
ADMIXTURE bar plot showing inferred ancestry proportions (K = 3) based on SNP data from brook stickleback populations in Astotin Lake, Muir Lake, and Shunda Lake. Each column represents one individual. Within lakes, individuals are ordered left-to-right by pelvic phenotype (spined, intermediate, unspined).

